## Supplementary figures and images for "Exploring the overlap between rheumatoid arthritis susceptibility loci and long non-coding RNA annotations"

### Supplementary figure 1

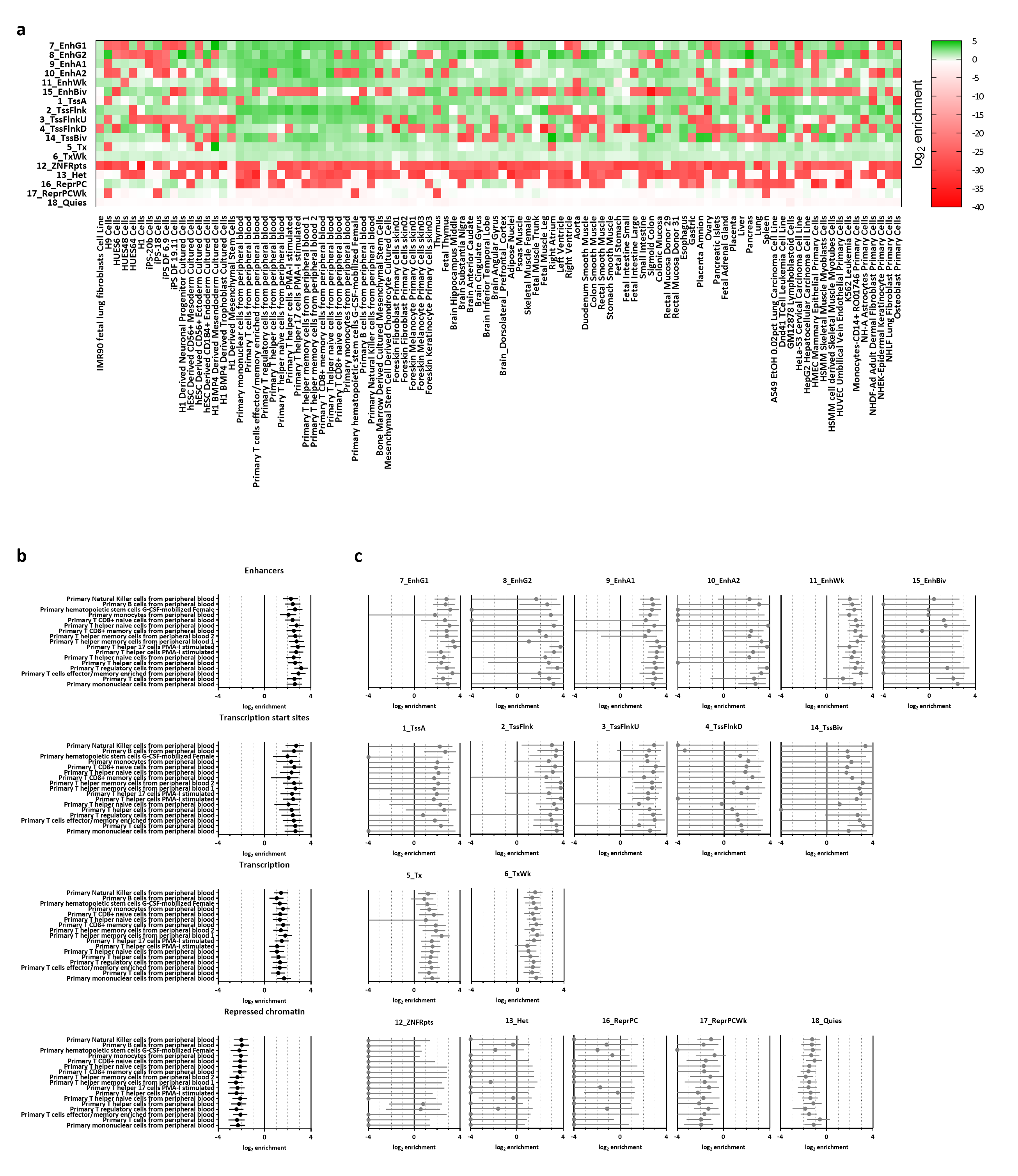

### Supplementary figure 2

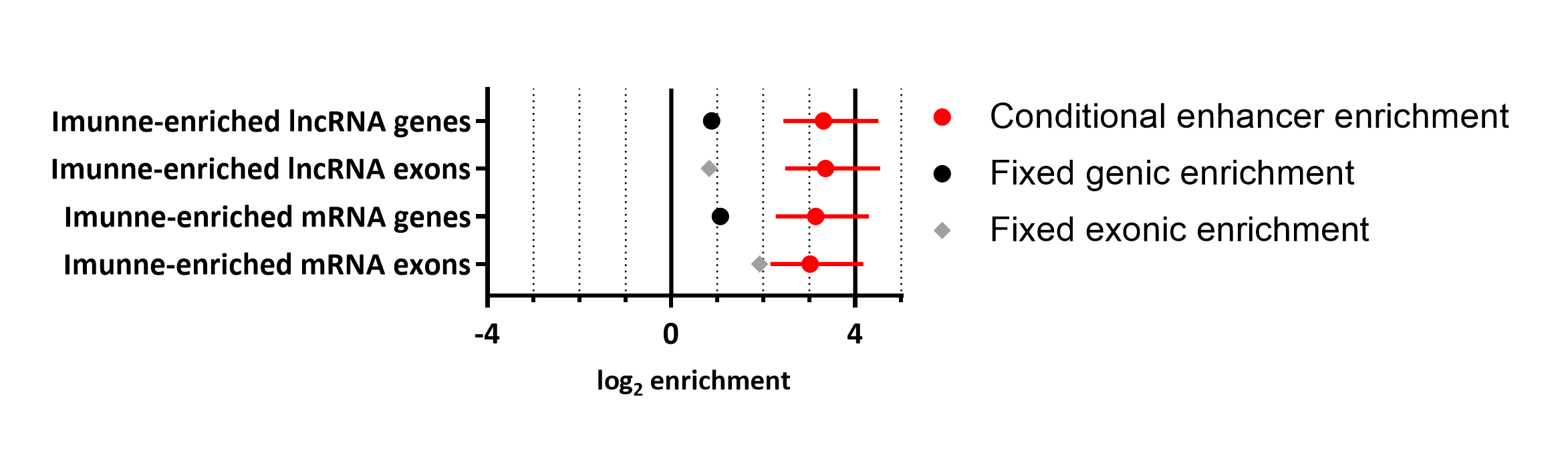
